# Durability of Motor Learning by Observing

**DOI:** 10.1101/2023.10.25.563597

**Authors:** Natalia Mangos, Christopher J. Forgaard, Paul L. Gribble

## Abstract

Information about another person’s movement kinematics obtained through visual observation activates brain regions involved in motor learning. Observation-related changes in these brain areas are associated with adaptive changes to feedforward neural control of muscle activation and behavioural improvements in limb movement control. However, little is known about the stability of these observation-related effects over time. Here we used force channel trials to probe changes in lateral force production at various time points (1 min, 10 min, 30 min, 60 min, 24 h) after participants either physically performed, or observed another individual performing upper limb reaching movements that were perturbed by novel, robot-generated forces (a velocity-dependent force-field). Observers learned to predictively generate directionally and temporally specific compensatory forces during reaching, consistent with the idea that they acquired an internal representation of the novel dynamics. Participants who physically practiced in the force-field showed adaptation that was detectable at all time points, with some decay detected after 24 h. Observation-related adaptation was less temporally stable in comparison, decaying slightly after 1 h and undetectable at 24 h. Observation induced less adaptation overall than physical practice, which could explain differences in temporal stability. Visually acquired representations of movement dynamics are retained and continue to influence behavior for at least one hour after observation.

**Significance:** We used force channel probes in an upper limb force-field reaching task in humans to compare the durability of learning-related changes that occured through visual observation to those after physical movement practice. Visually acquired representations of movement dynamics continued to influence behavior for at least one hour after observation. The significance of the findings is two-fold. First, they broaden our understanding of the potential role of visual observation in naturalistic motor skill acquisition. Our findings point to a (at least) 1-hour window during which visual observation of another person (e.g. a tutor or a conspecific learner) could play a role in motor learning, especially in early learning, presumably (as shown in our previous work) through activation of the neural circuitry that underlies motor skill acquisition. Second, this one-hour window represents an opportunity for observation-based interventions in neurorehabilitation (e.g. interleaving observation with physical practice) to aid in driving cortical reorganization for recovery of motor function.

## Introduction

Skilled action depends on the brain’s ability to acquire and modify neural representations of movement dynamics—the relationship between movement kinematics and the forces required to generate those movements. Learning about the forces required for movement is crucial for both controlling voluntary multi-joint movements such as upper limb reaching (Hollerbach and Flash, 1982; Koshland et al., 1991; Gribble and Ostry, 1999; Sainburg et al., 1999; Maeda et al., 2017), for using tools (Solomon et al., 1989), and for lifting objects (Flanagan and Wing, 1997). Regulation of the forces involved in movement is also a critical aspect of stroke recovery (Kokotilo et al., 2009), during which patients often require re-learning basic movement skills such as reaching. Skilled movement critically depends on predictive, feedforward neural representations of forces (see Wolpert et al., 2011). There is a large body of literature examining how physical movement practice drives this adaptive process. Recent work suggests that changes in neural representations of movement dynamics can also be acquired in the absence of movement, by observing the actions of another individual performing movements in novel dynamical environments (Mattar and Gribble, 2005; Wanda et al., 2013). While the temporal stability of motor learning obtained through physical movement practice has been well documented, the durability of motor learning by observing is unknown. This is the focus of the studies described here.

Evidence from force-field reaching tasks in both humans and non-human primates suggests that motor learning through physical practice results in the acquisition and use of a novel representation of reach dynamics (Shadmehr and Mussa-Ivaldi, 1994; Conditt et al., 1997; Li et al., 2001; Gandolfo et al., 2000; Gribble and Scott, 2002). Recent studies provide evidence that such an adaptive process also occurs in human observers who observe the movements of another individual learning to adapt to a novel force-field. Observers are able to predictively generate a novel, time-varying pattern of muscle forces that counteracts the force-field that perturbed the observed movements. This occurs despite never physically experiencing the force-field themselves (Mattar and Gribble, 2005). This effect does not seem to depend upon the use of conscious, explicit strategies for movement, but rather on the implicit engagement of the sensorimotor system (Mattar and Gribble, 2005). The findings of Mattar and Gribble (2005) have since been replicated a number of times (Brown et al., 2009, 2010; Williams and Gribble, 2012; Bernardi et al., 2013; McGregor and Gribble, 2015; McGregor et al., 2016, 2018a) and are consistent with those of Wanda et al. (2013), in which observation-induced force generation patterns were measured directly, using error clamp probe trials (Scheidt et al., 2000). Wanda et al. (2013) showed that as is the case with force-field learning through physical practice, observation of force-field learning allowed observers to generate novel patterns of muscle force that mirrored the timing and direction of the force-field.

Temporally and directionally specific changes in predictive limb control are detectable shortly after either observing force-field learning or physically practicing in a force-field; however, the extent to which adaptation can be retained, and influence future behaviour, has not been well described for motor learning by observing. Studies of force-field adaptation through physical practice provide evidence that its effects can persist past the end of the training period. For example, adaptive changes in predictive force output patterns can be retained for at least 24 hours following physical force-field exposure (Joiner and Smith, 2008), and adaptive changes in movement kinematics have been shown to persist when tested 5 months after adaptation (Shadmehr and Brashers-Krug, 1997). Physical force-field adaptation also produces savings—a phenomenon in which re-learning occurs at a faster rate than initial learning, even if after-effects of the initial learning have been washed out (Mathew et al., 2021; Shadmehr and Brashers-Krug, 1997). Physical force-field adaptation may therefore give rise to durable changes in the mechanisms underlying skilled action, improving future performance on the same task and on previously untrained but similar tasks (Conditt et al., 1997; Gandolfo et al., 1996; Goodbody and Wolpert, 1998; Huang and Shadmehr, 2007; Hwang et al., 2003; Malfait et al., 2002; Sainburg et al., 1999; Shadmehr and Moussavi, 2000).

To date, the temporal stability of the adaptation that occurs via observing force-field learning has not been characterized. Most data on the effects of observing force-field learning were obtained during or only minutes after observation had occurred (Brown et al., 2009, 2010; Malfait et al., 2010; Mattar and Gribble, 2005; McGregor et al., 2018a,b; McGregor and Gribble, 2017; Williams and Gribble, 2012). Two studies have been reported in which performance in a force-field (McGregor and Gribble, 2015) and on a somatosensory perception task (Bernardi et al., 2013) were assessed approximately one hour after observing. Investigating the durability of adaptation that occurs through observation is useful for two reasons. First, it could allow us to draw comparisons to the temporal stability of the learning that occurs through physically reaching in a force-field. If learning that occurs through observation and learning that occurs through physical practice were found to be similarly durable under the same conditions, this would be consistent with the idea that these two processes share common neural mechanisms. Second, a tool that can elicit temporally stable changes in the neural control of movement without physical practice could be useful in a clinical setting, particularly for promoting the recovery of motor function after stroke. In a clinical context, information about the durability of the learning that occurs through observation may be relevant to the design of observation-related approaches to neurorehabilitation.

We measured force generation patterns before and after human participants had either observed force-field learning or physically practiced reaching in a novel force-field. Following either observation or physical practice, we introduced a delay lasting between one minute and 24 hours before a second set of force measurements were obtained. We probed adaptation by measuring predictive force generation patterns after observational or physical force-field exposure and the variable-length delay, which allowed us to characterize and compare the temporal stability of the adaptation elicited by each type of force-field exposure. We also characterized the stability of the adaptation by comparing the rates at which changes in force output were washed out over consecutive reaches. Adaptation following observation of force-field learning continued to influence behavior for at least one hour after observation.

## Materials and Methods

### Participants

A total of 178 individuals participated in this study. Participants reported no history of neurological or musculoskeletal disorders. Eligibility requirements included self-reported right-handedness, no prior experience with a force-field learning paradigm, and no self-reported visual, neurological, or musculoskeletal disorders. Eighteen participants were excluded for failure to complete the testing session, failure to meet the originally stated eligibility criteria (disclosed after completing the testing session), failure to follow instructions during the testing session, or disruption of the testing session by malfunctioning equipment. New participants were recruited until each experimental group contained 16 participants (after exclusions), for a total of 160 participants (mean age: 20 yr± 4 yr). All study procedures were approved by Western University’s Research Ethics Board.

### Experimental Setup

Participants grasped the handle of a two-joint robotic manipulandum (KINARM Laboratories, Kingston, Canada) with their right hand and performed planar reaching movements to eight visual targets. A custom airsled was placed beneath the right arm to reduce fatigue from supporting the arm against the gravitational load. Targets were displayed on a semisilvered mirror mounted horizontally between eye-level and the hand. This surface occluded participants’ direct vision of the handle and their right arm. The position of the handle was displayed as a circular white cursor. In addition, a transient tracing of the handle’s recent trajectory was displayed behind the cursor (shown as a red trace in Fig. 1). Targets were spaced equally around the circumference of a circle with a 10 cm radius. A home target was located at the center of the circle. On a given trial, participants moved the robot handle to the home target until prompted, by the appearance of one of the eight possible movement targets, to perform a straight line reaching movement to that target. Shortly after the movement target was reached, the robot moved the participant’s passive arm so that the handle was placed back at the home target. Movement targets appeared in random order within bins of eight trials, such that each target appeared once per bin.

**Figure 1:**
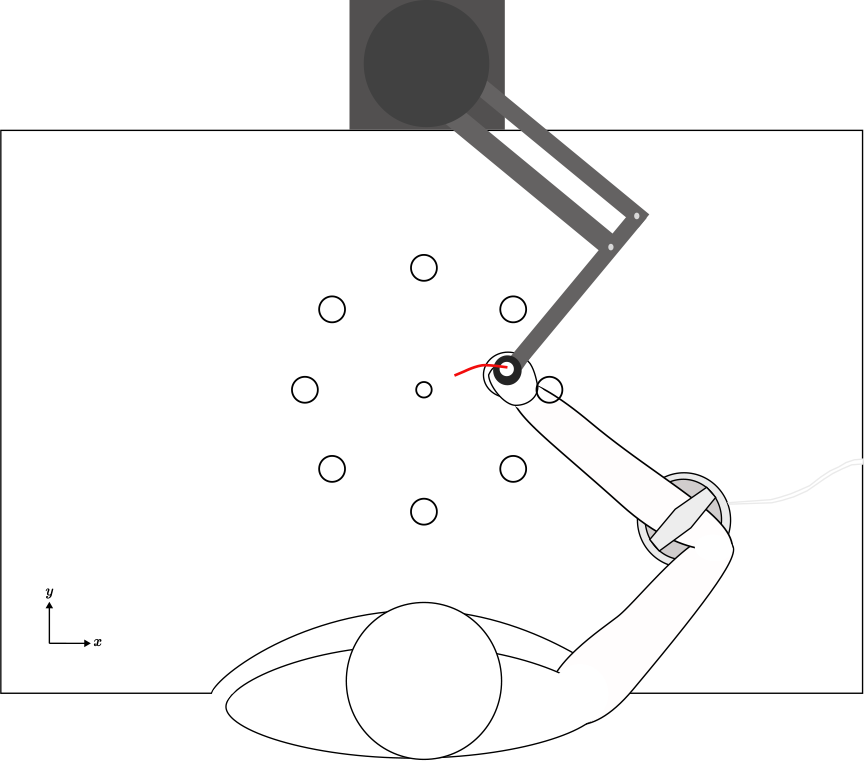
Experimental Setup. Participants were seated in front of a two-joint robotic manipulandum. A custom air-sled was placed on top of the table to support the arm. Direct vision of the arm was completely occluded by the opaque, horizontal display (depicted here as translucent) onto which targets were projected.

To regulate reach velocity, participants received color feedback after each reach indicating whether the reach was completed too slowly (target turned blue), too quickly (red), or within the desired time window of 400 ms ± 50 ms (target disappeared rather than changing color). Time constraints did not include reaction time; participants were told that the color feedback would reflect the amount of time elapsed between leaving the home target and reaching the movement target. Participants were instructed to reach to the target in a straight line and to do so within the correct time constraints, to stop on the target rather than reaching through it, and to wait at the target position until the robot initiated a return back to the home target.

### Task Design

The experiment was divided into four blocks (Fig. 2 A). Participants were assigned to one of 10 experimental groups (16 participants per group). Each group designation referred to which of two possible force-field learning protocols (movement or observation) and which of five possible delay period protocols (1 min, 10 min, 30 min, 60 min, or 24 h delay) a participant completed.

**Figure 2:**
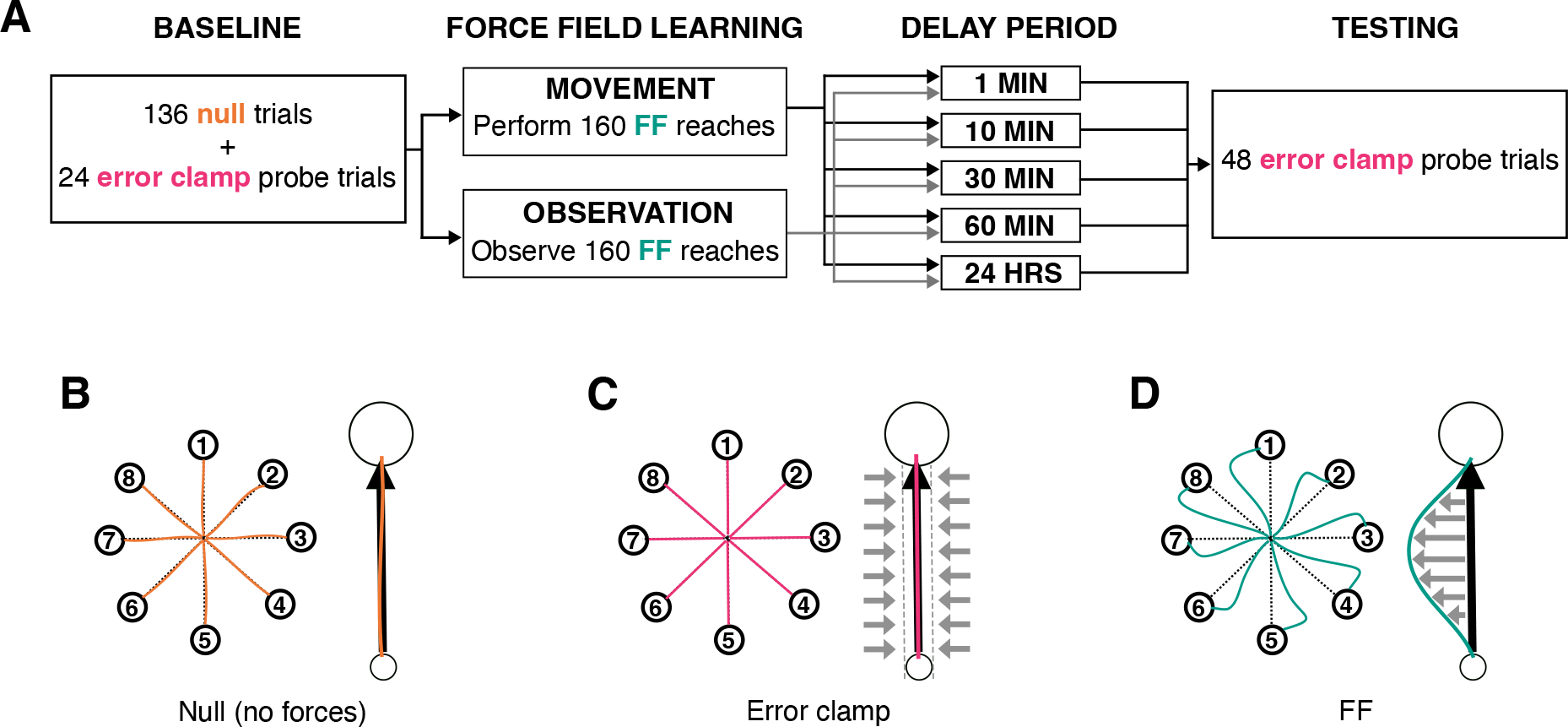
Task design. A. Experimental blocks. Ten groups of participants completed a Baseline block, force-field (FF) Learning block, Delay Period block, and Testing block. All participants completed the same Baseline and Testing blocks; which FF Learning protocol (movement or observation) and Delay Period protocol (1 min, 10 min, 30 min, 60 min, or 24 h delay) participants completed was manipulated. Force generation patterns were probed during error clamp trials in baseline and testing blocks. B–D. Hand paths during null trials, error clamp trials, and early FF trials—for reaches to all eight targets (left) or for one reach to an individual target (right). The large black arrow indicates the intended reach direction; small grey arrows depict robot-imposed forces.

### Baseline block

Participants began by performing 160 reaches in the absence of any forces applied by the robot (null environment; Fig. 2 B). The latter half of trials in this block included 24 pseudorandomly interspersed error clamp trials (Scheidt et al., 2000)—probe trials in which the robot restricted the position of the handle to a straight line channel between the home target and the end-target (Fig. 2 C). Such trials allowed us to quantify participants’ lateral force output against the channel walls over the time course of a reach, prior to any force-field exposure.

### Force-field learning block

Following the baseline block, participants completed one of two possible force-field learning protocols. Participants assigned to a movement group performed 160 reaches in a counter-clockwise velocity-dependent force-field environment. The forces applied by the robot were defined by Equation 1:

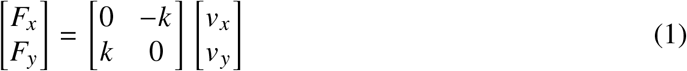

where *F_x_* and *F_y_* are robot forces, *v_x_* and *v_y_* are hand velocities in the x– and y–axes (left-right and forwards-backwards in a horizontal plane) respectively, and *k* = 14 N s m^−1^ (Fig. 2 D). Participants assigned to an observation group did not perform reaches in a force-field environment; rather, they watched a video of a tutor performing 160 reaches in the counter-clockwise force-field environment given by Equation 1. The video depicted a top-down view of the tutor’s right arm as they learned to make straight movements in the presence of perturbing forces (Fig. 3).

**Figure 3:**
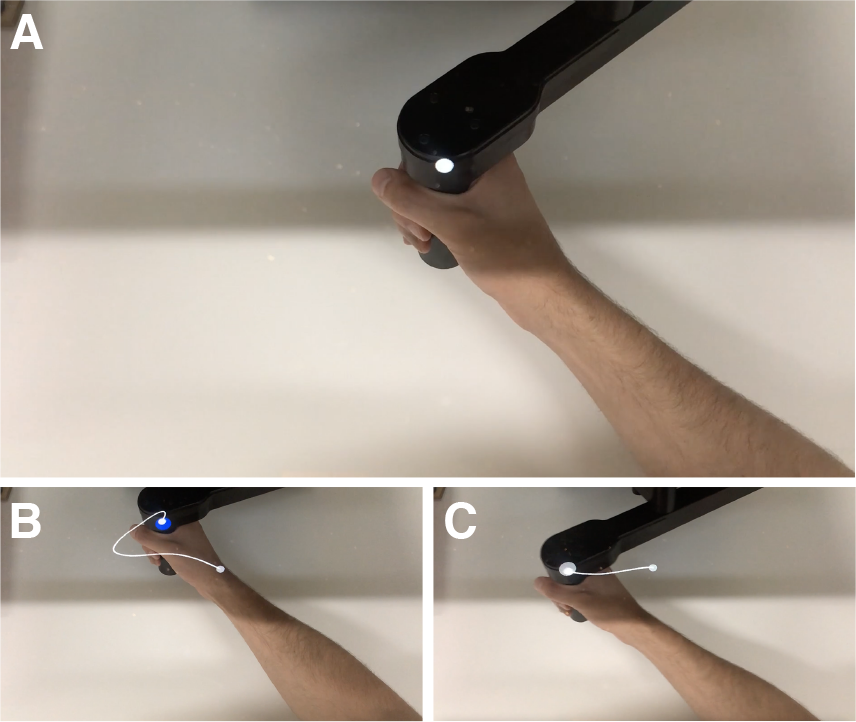
Sample frames from the video used for observation groups. A. Tutor resting at the home position. B. Reach completed during the early stages of force-field adaptation. The tutor experiences reaching errors (large trajectory curvature). C. Reach completed during the late stages of force-field adaptation.

Participants assigned to an observation group remained seated in front of the robot while the video was projected onto the horizontal display. The video provided a first-person view of the tutor’s arm, from a perspective that would closely match a participant’s view of their own arm if it was visible when physically performing reaching movements using the robot. Participants were not informed of the presence of robot-generated forces in the video they observed. They were only told that the video showed someone performing reaches. Observers were instructed to keep count of the number of times the actor in the video reached the movement target in the desired time window, by counting the number of times the movement target disappeared rather than changing color. To verify that observers were paying attention during the video, they were asked to report their counts verbally, to be documented by the researcher, at each of four checkpoints throughout the video. Any participant who reported a number not within 20 % of the correct value at any of the four checkpoints was excluded from data analysis. Only one participant—who disclosed that they had fallen asleep during the video—reported a count that fell outside of this range and was dismissed with full compensation (and their data excluded from the study). Because producing voluntary movements while observing force-field learning has been shown to interfere with the effects of observing (Mattar and Gribble, 2005) participants were instructed to keep their hands resting flat on the table beneath the display, and to remain as still as possible throughout the duration of the video.

### Delay Period

After the force-field learning block, participants completed one of five possible delay periods: 1 min, 10 min, 30 min, 60 min, or 24 h. During delay periods of 60 min or shorter, participants remained seated in the robot chair and kept their hands resting flat on the table. Because we were unsure of whether moving during the delay period would retroactively interfere with the effects of learning by observing, participants were instructed to remain as still as possible throughout the duration of the delay period and not to move their arms. The researcher was present to monitor participants. Any movement observed by the researcher was accompanied by a verbal reminder that participants should stay as still as possible and not move their arms. Participants were permitted to listen to a podcast or speak with the researcher during the delay period, and throughout the delay the horizontal display showed only a black screen.

Participants assigned to a group with a 24 h delay left the testing facility immediately following completion of the force-field learning block and returned 24 h ± 2 h later.

### Testing Block

Following the delay, participants were instructed to perform a final set of reaches with the same rules as in the baseline block. The testing block consisted of 48 error clamp trials, which allowed us to obtain measurements of participants’ lateral force output following the delay period.

### Analyses

The force-field used in this experiment perturbed the arm laterally relative to the reach direction (Fig. 2). Adaptation could therefore be measured as the change in lateral forces generated by participants during reaching. For each error clamp trial in the baseline and testing blocks, the forces that restricted the hand’s trajectory to a straight line were generated by the robot, which produced forces that mirrored the lateral forces generated by the participant. Accordingly, measurements of participants’ lateral force output over the time course of a given reach were obtained by taking the sign-flipped time series of robot-generated forces. All kinematic and force data were digitally sampled at 1000 Hz and low-pass filtered at 10 Hz using a double-pass, third-order Butterworth filter implemented in MATLAB. Time-series force and velocity data were aligned on peak tangential hand velocity and a window of data from 400 ms before to 400 ms after the time of the peak was extracted. Reaches were excluded in cases in which a participant sped up and slowed down multiple times between leaving the home target and reaching the movement target, as determined by the presence of two or more acceleration phases separated by a deceleration phase. Fewer than 4 % of trials were excluded for this reason.

### Adapted and ideal lateral force profiles

For each participant, lateral force data from error clamp trials in the baseline block were collapsed across same-target trials to generate one average baseline force output profile per target. Then, for each trial in the testing block, an adapted lateral force output profile was generated by subtracting the target-matched baseline time series from the lateral force profile for that testing block trial. Adapted lateral force profiles therefore represented the change, from pre-to post-force-field exposure, in the magnitude and temporal pattern of lateral forces produced by participants during reaching. For each trial in the testing block, we also generated an ideal lateral force output profile, which gave the magnitude and temporal pattern of lateral forces that would have been required to perfectly oppose the force-field if the force-field had been applied during that trial. The ideal profile for a given reach was computed according to Equation 1, taking as input the instantaneous velocity of the hand over the time course of the reach. Because the force-field was designed to perturb the arm counter-clockwise relative to the direction of the reach, compensatory forces produced by participants were those exerted in the clockwise direction. Accordingly, for simplicity we re-signed all lateral force data to make the sign convention for clockwise lateral forces positive.

### Fraction of ideal lateral force exerted

We quantified adaptation as the fraction of required compensatory forces that participants learned to produce after force-field exposure and the delay period. For each trial in the testing block, we computed the fraction of ideal force exerted by integrating over time a participant’s adapted and ideal lateral force profiles, and then expressing the former integral as a fraction of the latter. We took the average fraction of ideal force exerted over the first eight reaches of the testing block (one to each target) as our primary quantitative measure of adaptation, and assessed group differences in adaptation using a two-way analysis of variance (ANOVA) with type of force-field exposure (movement, observation) and delay length (1 min, 10 min, 30 min, 60 min, or 24 h) as factors.

### Washout rate

Linear regression of the fraction of ideal force exerted onto trial number was used to determine the rate at which the adaptation washed out over successive testing block trials. A washout rate for each participant was given by the slope of the line that best fit (in a least-squares sense) the fraction of ideal force exerted per trial over the 48 trials in the testing block. Washout rates were compared between groups via two-way ANOVA, with type of force-field exposure (movement, observation) and delay length (1 min, 10 min, 30 min, 60 min, or 24 h) as factors.

### Velocity dependence

Because during force-field reaching the magnitude of robot-imposed perturbing forces scaled with reach velocity, we also assessed the extent to which participants’ adapted forces correlated with reach velocity. For each participant, we computed Pearson’s *r* for the correlation between the average adapted force and velocity profiles over the first eight reaches of the testing block. We performed multiple one-tailed t-tests to assess whether there was a significant positive correlation between adapted force output and reach velocity in each experimental group. Bonferroni-Holm corrections were applied for two families of five delay lengths (1 min, 10 min, 30 min, 60 min, or 24 h).

### Statistical analyses

ANOVAs and associated post-hoc tests were carried out in JASP v0.17.1. Where assumptions of homoscedasticity were violated (Levene’s *p* < 0.05), ANOVA models were adjusted using the weighted least squares method (i.e., observations were reweighted proportionally to the reciprocal of the error variance). All other computations and statistical analyses were performed in MATLAB R2021b.

## Results

### Adaptation of force output

Participants were assigned to one of 10 experimental groups, each involving one of two types of force-field exposure (movement or observation) immediately followed by one of five possible delay lengths (1 min, 10 min, 30 min, 60 min, or 24 h). All participants performed error-clamped reaches before (baseline block) and after (testing block) force-field exposure and the delay period, providing us with direct measurements of the lateral forces they generated during those reaches.

Figure 4 A–B (left panel) shows the average lateral forces generated across baseline block reaches for two experimental groups—the movement and observation groups that had the shortest delay length (1 min). Note that force profiles are aligned on peak velocity at 0 ms, and positive values represent forces generated in the direction opposite the force-field (i.e., clockwise). As expected, participants in both observation and movement 1 min groups produced minimal lateral forces on reaches performed prior to any force-field exposure. The center column shows average lateral forces across the first eight reaches of the testing block (one reach to each target). These force profiles reflect the earliest measurements of participants’ force production patterns following the delay period. Both the group that observed and the group that physically interacted with the force-field produced patterns of forces that were clearly different from what they produced in the baseline block.

**Figure 4:**
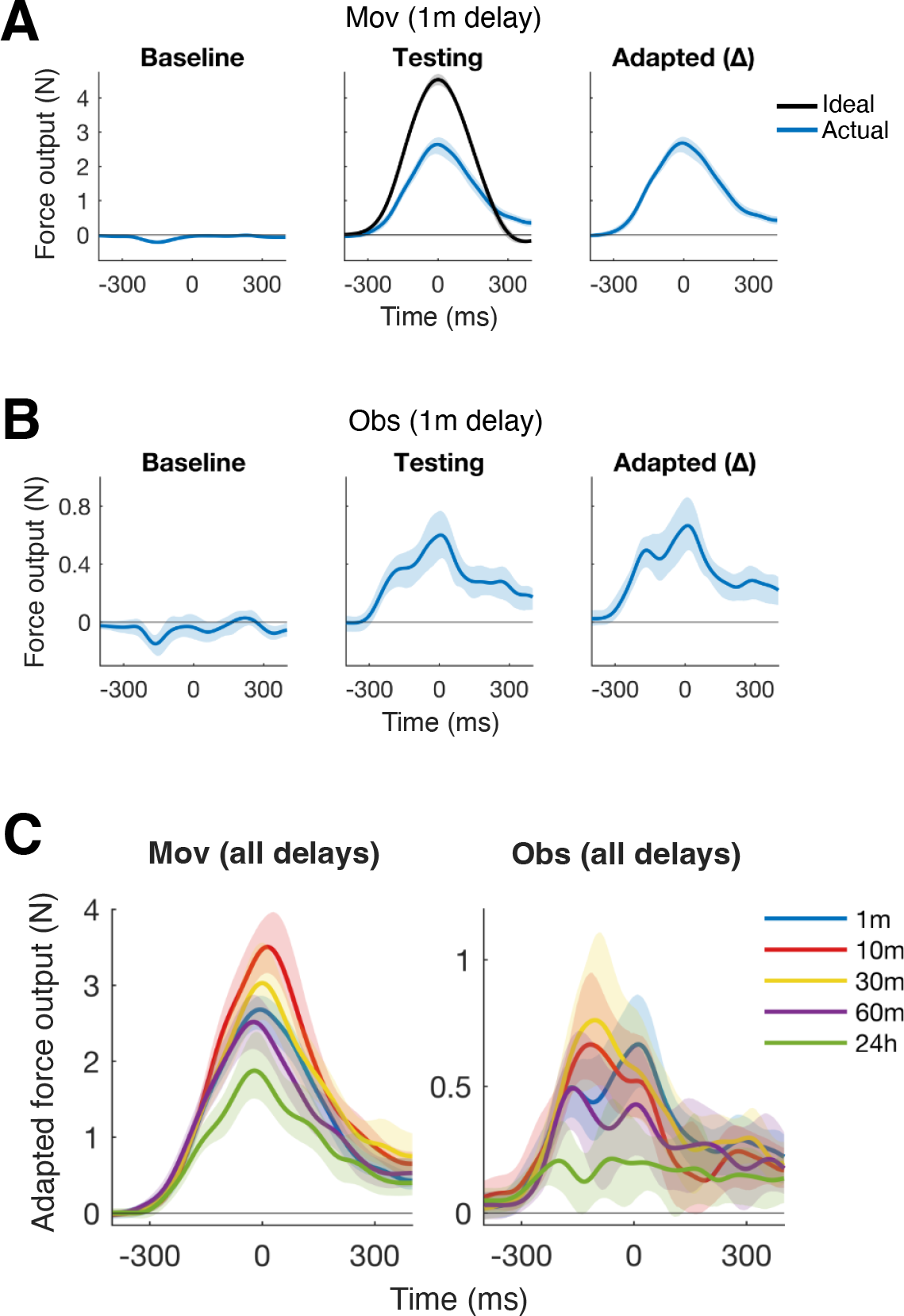
Adaptation of lateral force output after force-field exposure (movement or observation) and a variable delay period. A and B: Average lateral forces generated during baseline block reaches (left) and during the first eight reaches of the testing block (center), across participants in the movement (1 min delay, A) or observation (1 min delay, B) groups. The black trace in the center panel of A is the average lateral force needed to perfectly counteract the force-field (i.e., the ideal lateral force), computed from the velocities of the first eight testing block reaches. Adapted force profiles (right) are calculated as the average per-target change in lateral force from baseline to the first eight reaches of the testing block. Shaded regions are 95 % CIs across participants, using 1000 bootstraps. Force profiles are centered on peak velocity at 0 ms, and positive y-values represent forces generated in the compensatory (clockwise) direction. C: Adapted force profiles, averaged over the first eight reaches of the testing block, for movement (left) and observation (right) groups with different delays between force-field exposure and testing.

Participants who physically practiced reaching in the force-field and were tested after 1 min generated lateral forces that rose to their peak around the time of maximum reach velocity and then fell again, creating an approximately bell-shaped force profile that matched the direction and temporal pattern of forces required to oppose the force-field (Fig. 4 A, center panel). The lateral forces produced by participants who observed force-field learning also followed this pattern (Fig. 4 B, center panel), although they were lesser in magnitude and more variable. The change in force output relative to baseline is shown in the rightmost column of Fig. 4 A–B, which depicts the average per-target change in lateral forces relative to baseline across the first eight reaches of the testing block.

For the other eight experimental groups (movement or observation with delays ranging from 10 min to 24 h), the lateral force profiles bore some similarities to those depicted in Figure 4 A–B. Baseline force output profiles for all experimental groups closely resembled those depicted in Fig. 4 A–B, where participants generated near-zero lateral forces but for a very slight tendency to lean counter-clockwise on average. Following force-field exposure and the delay period, participants in all groups adapted their lateral forces to push in the clockwise direction instead; however, the extent of adaptation varied between groups. This can be seen in Figure 4 C, which shows for each experimental group the average adapted lateral force profile across the first eight reaches of the testing block. Participants who physically reached in the force-field showed clear evidence of adaptation after all delay lengths (Fig. 4 C, left panel)—and despite some decay after 24 h, adapted lateral forces for the five movement groups generally held the same shape. This was not the case for participants who observed force-field learning. The right panel of Figure 4 C shows that adapted lateral forces for observers following delays of 1 min, 10 min, and 30 min were similar in magnitude and held a bell-like shape—but after a 60 min delay, observers’ lateral force profiles were more variable. After 24 h observers produced only a faint and variable signal of adaptation.

For each experimental group, we quantified the amount of adaptation that occurred as the average fraction of ideal force exerted across the first eight reaches of the testing block (Fig. 5 A). Group differences were assessed by two-way ANOVA, with type of force-field exposure (movement, observation) and delay length (1 min, 10 min, 30 min, 60 min, or 24 h) as factors. We found a significant main effect of type of force-field exposure *F* (1, 150) = 337.79, *p* < 0.001 and a significant main effect of delay length *F* (4, 150) = 8.68, *p* < 0.001 on the fraction of ideal force exerted; however, these effects were qualified by a significant interaction effect *F* (4, 150) = 3.32, *p* = 0.012. A simple main effects analysis showed that type of force-field exposure affected the fraction of ideal force exerted after every delay length (all *p* < 0.001), and that delay length affected the fraction of ideal force exerted after physical force-field exposure (*p* < 0.001) and after observation (*p* = 0.019).

**Figure 5:**
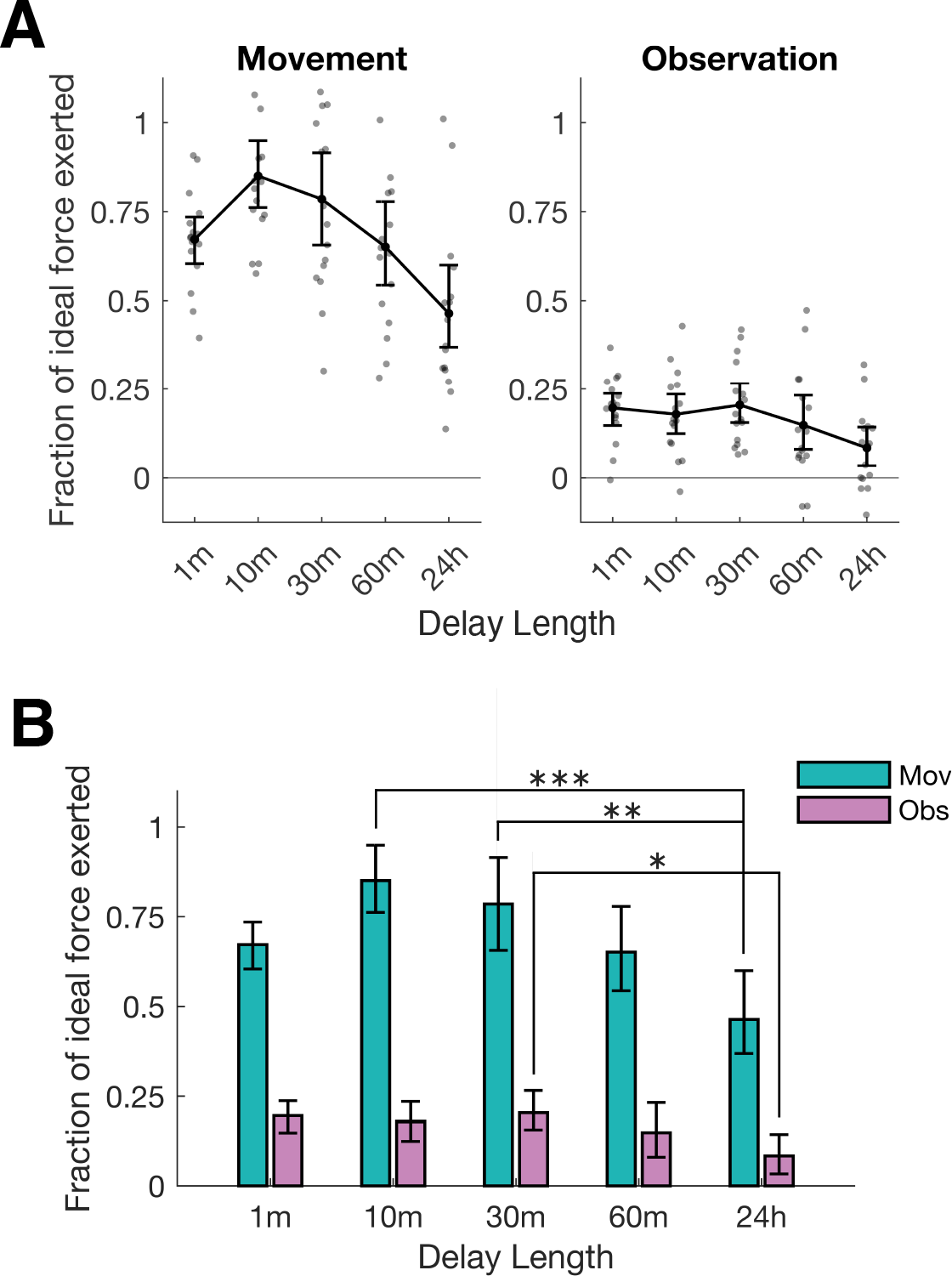
Quantity of adaptation after force-field exposure (movement or observation) and a variable delay period. A: Amount of adaptation, quantified as the average fraction of ideal lateral force exerted across the first eight testing block trials. Values equal to 1 reflect perfect adaptation and values equal to 0 reflect no adaptation. Means and 95 % CIs, from 1000 bootstraps, are shown for movement (left) and observation (right) groups with different delays. Closed circles represent individual participant means. B: Mean (±95 % bootstrap CIs) amount of adaptation across delay lengths, for movement and observation groups. All values of mean adaptation for participants in the movement group are significantly different from each mean for the observation groups (*p* < 0.001). Significant differences between groups with the same type of force-field exposure but different delay lengths are depicted by asterisks: ^*^ *p* < 0.05, ^**^ *p* < 0.01, ^***^ *p* < 0.001.

Tukey-corrected post-hoc tests were conducted to assess differences in adaptation between groups with the same type of force-field exposure but different delay lengths. Pairwise comparisons were carried out using 1000 bootstraps, and *p*-values were corrected for families of five tests. Within the first 60 min after either type of force-field exposure (movement or observation), mean adaptation did not differ significantly between groups that experienced different delay lengths (all *p* > 0.05). Among participants who physically reached in the force-field, those who experienced a 24 h delay before the testing block produced a significantly lower fraction of ideal force than participants whose delay lasted only 10 min (*p* < 0.001) or 30 min (*p* = 0.001; Fig. 5 B). Participants who observed force-field learning and then experienced a 24 h delay produced a significantly lower fraction of ideal force than observers who waited 30 min before starting the testing block (*p* = 0.049; Fig. 5 B).

### Washout rate

Next, we assessed the effect of delay length on the rate at which reaching-or observation-related adaptation washed out during error clamp probe trials after the delay period. Figure 6 A shows the mean fraction of ideal force exerted on each trial in the testing block, for each of the 10 experimental groups. Effects generally washed out more slowly after observing than after physically reaching in the force-field. Washout rates appeared similar between all observation groups, as well as between movement groups with delays lasting between 1 min and 60 min; however, effects appeared to wash out more slowly in the movement group that experienced a 24 h delay than in the other movement groups (Fig. 6 A).

**Figure 6:**
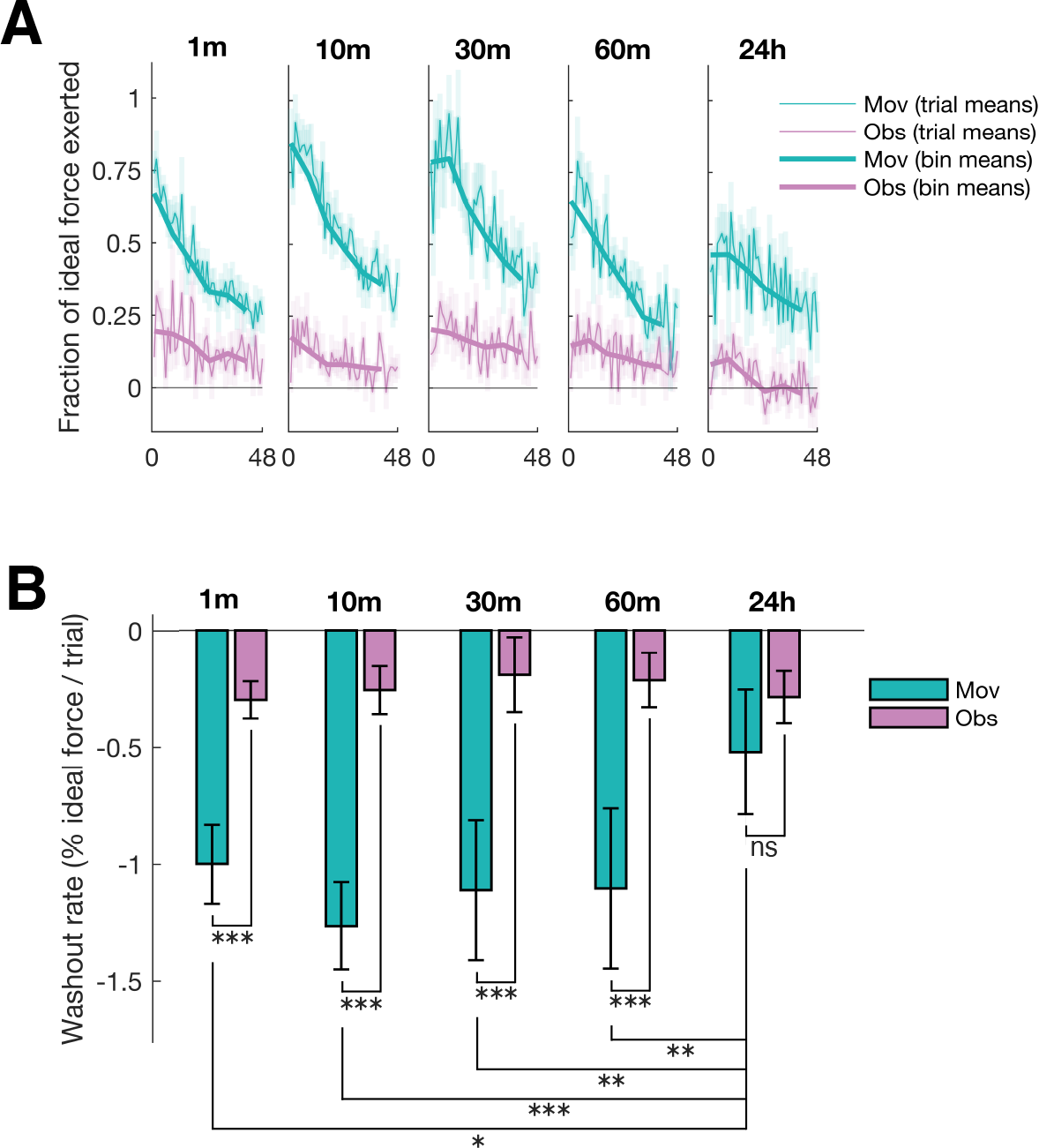
Washout of adaptation during error clamp trials. A: Fraction of ideal force exerted across all 48 testing block trials, for each group. Thin lines and surrounding shaded regions represent single trial means ± SE across participants in a given group; thick lines represent means for bins of eight trials (one reach to each target per bin). B: Mean washout rates (±95 % CIs, using 1000 bootstraps). Washout rates for individual participants were computed as the slope of the line that best fit the fraction of ideal force exerted across the 48 testing block trials. Washout rates were faster for movement than observation groups after all delays except 24 h. Washout rates did not differ significantly between observation groups. Washout rates were faster for movement groups with delays lasting 1 min–60 min than for the movement group with a 24 h delay. ^*^ *p* < 0.05, ^**^ *p* < 0.01, ^***^ *p* < 0.001, ns *p* > 0.05.

For each participant, the fraction of ideal force generated across the 48 testing block trials was modeled as a linear function of trial number, where the rate of washout was given by the slope of the least-squares line of best fit. Average washout rates were compared between groups by two-way ANOVA, with type of force-field exposure (movement, observation) and delay length (1 min, 10 min, 30 min, 60 min, 24 h) as factors. There were significant main effects of type of force-field exposure

*F* (1, 150) = 156.40, *p* < 0.001 and delay length *F* (4, 150) = 4.56, *p* = 0.002 on washout rate; however, these effects were qualified by a significant interaction *F* (4, 150) = 5.84, *p* < 0.001. A simple main effects analysis showed that delay length affected washout rate after physical force-field exposure (*p* < 0.001) but not after observation (*p* = 0.589), and that type of force-field exposure affected washout rate after every delay length (*p* < 0.001) except 24 h (*p* = 0.086; Fig. 6 B). Since group-level washout curves appeared similar between movement groups with delays of 60 min or less (Fig. 6 A), we compared the washout rate of each movement group to that of the 24 h group via Dunnett’s test. We found that effects washed out significantly slower 24 h after physical force-field exposure than after 1 min (*p* = 0.025), 10 min (*p* < 0.001), 30 min (*p* = 0.004), or 60 min (*p* = 0.004; Fig. 6 B).

### Velocity dependence

To characterize the correlation between each participant’s average adapted force and velocity profiles over the first eight reaches of the Testing block, we computed Pearson’s *r*. Group mean profiles and correlation coefficients are shown in Figure 7. Participants who physically reached in the force-field produced forces over time that correlated strongly with reach velocity after all delay lengths (multiple one-tailed t-tests, *r*≥0.87 and *p* < 0.001 for all movement groups). For observers, adapted forces correlated moderately with reach velocity within the first 30 min after force-field exposure (*r*≥0.48 and *p* < 0.001 for all observation groups). The association was weaker after 60 min (*r* = 0.30, *p* = 0.012) and was no longer significant after 24 h (*r* = 0.12, *p* = 0.117). Note that *p*-values presented here were corrected by the Bonferroni-Holm method, applied for two families of five delay lengths.

**Figure 7:**
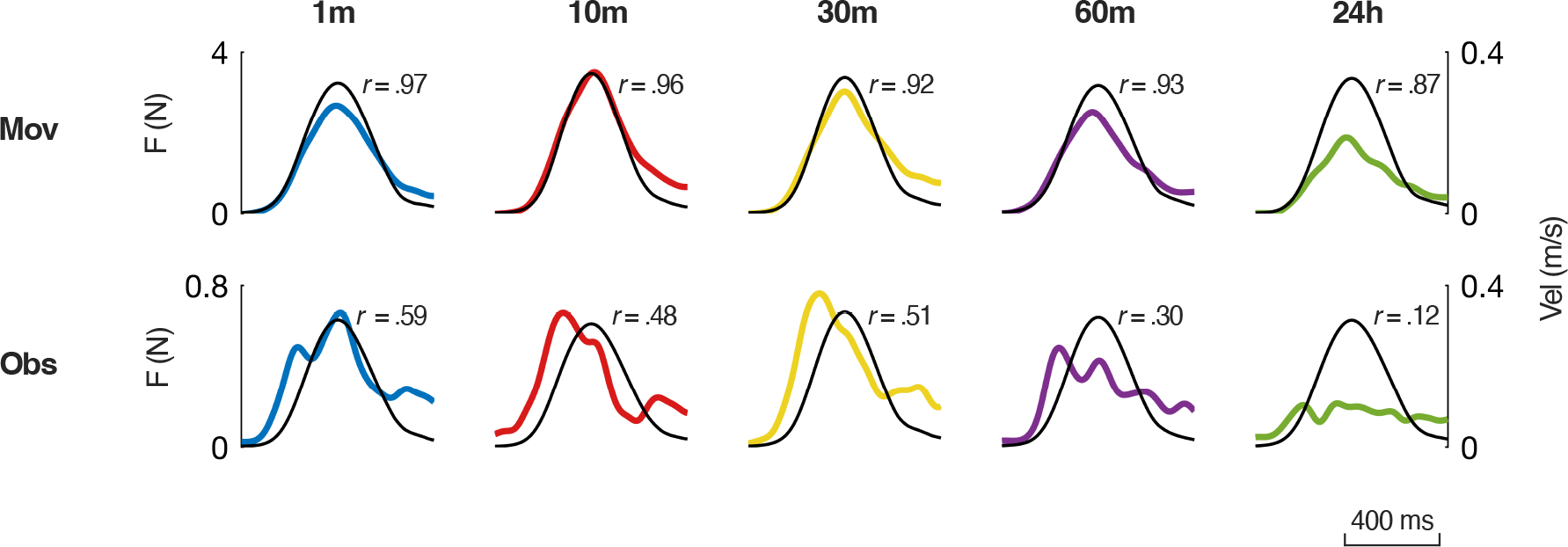
Velocity dependence of adapted force output. Each panel shows the adapted force profile (colored tracing, left axis) and velocity profile (black tracing, right axis) averaged across the first eight reaches of the testing block, for each group. Pearson’s *R* values shown here are group average correlation coefficients, where *R* was computed for the correlation between each participant’s average adapted force and velocity profiles over the first eight reaches of the testing block. Participants who physically reached in the force-field (top five panels) produced forces that correlated strongly with reach velocity after all delay lengths (multiple Bonferroni-Holm corrected one-tailed t-tests, all *p* < 0.001). For observers (bottom five panels), adapted forces correlated moderately with reach velocity within the first 30 min after force-field exposure (all *p* < 0.001). The association was weaker after 60 min (*p* = 0.012) and was no longer significant after 24 h (*p* = 0.117).

## Controls

### Differences in reach velocity

To determine whether there were group-level differences in movement speed during the first eight reaches of the testing block, we examined the mean peak reach velocity during these trials (Fig. 8). Differences in average peak reach velocity were assessed by two-way ANOVA. There was no significant main effect of delay length, *F* (4, 150) = 1.85, *p* = 0.122. There was a statistically reliable effect of type of force-field exposure (movement vs observation), *F* (1, 150) = 9.71, *p* = 0.002. There was also a statistically reliable interaction effect between delay length and type of force-field exposure, *F* (4, 150) = 3.18, *p* = 0.016. Bonferroni-corrected post-hoc tests returned non-significant (*p* > 0.05) results for all but one pairwise comparison, which found mean peak velocity to be reliably higher for the movement group with the 10 min delay than for the observation group with the 10 min delay (*p* = 0.002).

**Figure 8:**
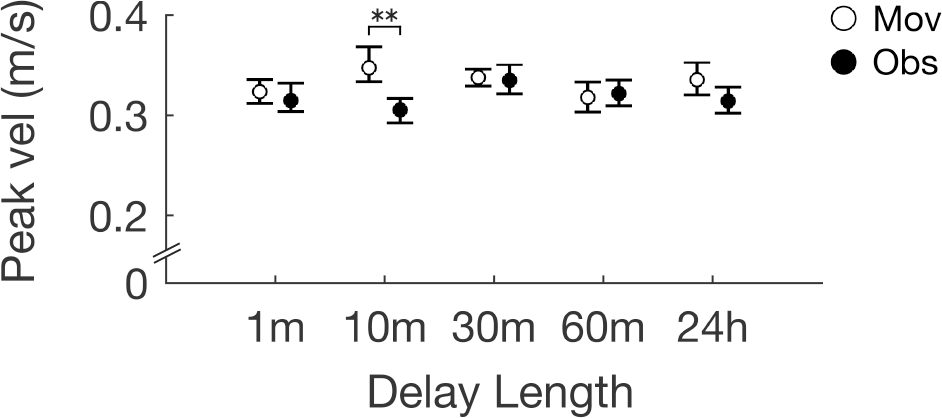
Mean peak reach velocity during the first eight reaches of the testing block. Open circles represent movement group means, and closed circles represent observation group means. Differences in average peak reach velocity between groups were assessed by two-way ANOVA with Bonferroni-corrected post-hoc tests conducted for all possible pairwise comparisons. The only statistically reliable difference found was that mean peak velocity was higher for the movement group with the 10 min delay than for the observation group with the 10 min delay (*p* = 0.002). All other comparisons were not significant (*p* > 0.05).

Despite the one difference noted above, peak velocity in the first eight trials of the testing block was not meaningful as a covariate when included in the ANOVA model we used to assess group-level differences in adaptation (*F* (1, 149) = 0.09, *p* = 0.765).

### 24-hour control group

We found small changes from baseline levels of lateral force among observers who experienced a 24 h delay between force-field exposure and the testing block (Figs. 4 C and 5); however, force profiles for this group of observers were rather variable and not meaningfully related to reach velocity (Fig. 7). This led us to suspect that the lateral forces produced by observers after a 24 h delay may not provide reliable evidence of observation-related adaptation. It could be that these forces are different on day two for reasons unrelated to force-field observation, such as changes in participants’ physical setup, behavioural changes associated with the absence of any practice or warm-up prior to beginning the testing block on day two, or other potential influence of leaving the facility and returning a day later.

To test for the possibility that lateral forces produced by observers after a 24 h delay may be present due to factors unrelated to force-field observation, we tested an additional control group of 16 participants (mean age 18.13 yr ± 0.34 yr) to complete a modified version of the 24 h condition of the experiment. The task was identical to the protocol completed by the observers who experienced a 24 h delay, but for one difference: participants in the 24 h control group did not complete a force-field observation block. Instead, control participants left the facility immediately after completing the baseline block, then returned 24 h ± 2 h later to complete the testing block.

Figure 9 A shows average lateral forces generated in the baseline block and across the first eight reaches of the testing block, for both participants who observed force-field adaptation followed by a 24 h delay, and for control participants who did not observe force-field adaptation. An adapted lateral force profile, averaged over the first eight reaches of the testing block, is also shown for each group. Lateral force profiles for the control group looked very similar to those measured for observers.

**Figure 9:**
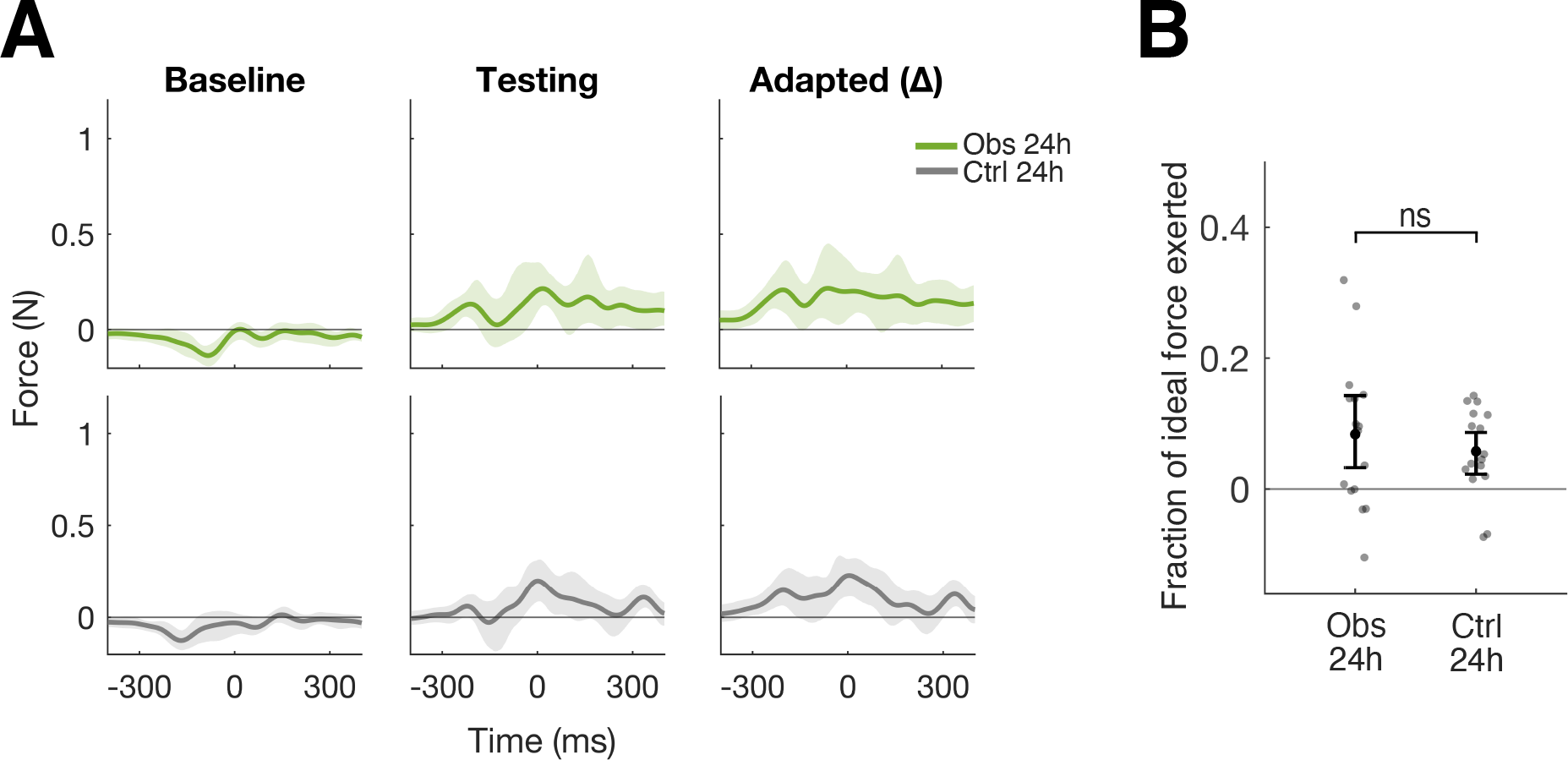
Results from the 24 h control group, compared to results from participants who observed force-field adaptation and were tested 24 h later. A: Average lateral forces generated during the baseline block reaches (left) and during the first eight reaches of the testing block (center), across observers (Obs 24 h, green) and the 24 h control group (Ctrl 24 h, grey). Adapted lateral force profiles (right) are calculated as the average per-target change in lateral force from baseline to the first eight reaches of the testing block. Shaded regions are 95 % CIs across participants, using 1000 bootstraps. Lateral force profiles are centered on peak velocity at 0 ms, and positive *y*-values represent forces generated in the compensatory (clockwise) direction. B: Amount of adaptation, quantified as the average fraction of ideal lateral force exerted across the first eight testing block trials, for observers and for the 24 h control group. Values equal to 1 reflect perfect adaptation and values equal to 0 reflect no adaptation. Means and 95 % CIs using 1000 bootstraps are shown for both groups, where closed circles represent individual participant means. There was no significant difference in the fraction of ideal lateral force exerted by observers who were tested after 24 h and the control group who did not observe force-field adaptation (randomization test, *p* = 0.228), suggesting that changes in lateral force seen 24 h post-observation do not provide reliable evidence of adaptation related to force-field exposure.

We quantified adaptation as the average fraction of ideal lateral force exerted across the first eight reaches of the testing block (Fig. 9 B), and we performed a randomization test using 10^4^ iterations to determine whether adaptation was reliably different between groups. We found no statistically reliable difference between the average fraction of ideal lateral force exerted by participants who observed force-field learning and were tested 24 h later and the control participants who did not observe force-field adaptation (*p* = 0.228). This makes it difficult to infer that the non-zero lateral forces produced by participants in the main experiment who observed force-field learning and were tested after 24 h are due to observation-related learning.

## Discussion

We characterized the durability of the learning that occurs through observing another individual undergoing force-field adaptation. We measured lateral forces generated by participants at 1 min, 10 min, 30 min, 60 min, or 24 h after either physically reaching in a force-field or observing another individual learning to reach in a force-field.

### Temporal Stability and Resistance to Washout

We found that like participants who learned to reach in a force-field through physical practice, observers also learned to predictively generate directionally and temporally appropriate compensatory forces during reaching. This finding is consistent with those previously reported by Wanda et al. (2013) and provides further support for the idea that neural representations of dynamics can be acquired by visual observation of movement (Mattar and Gribble, 2005).

Observing another individual undergoing force-field learning induced an adaptation of predictive limb control that showed no evidence of decay for at least 30 min after the end of the observation period. We also found reliable evidence of adaptation at 60 min post-observation, although temporal force profiles in this case were less closely related to reach velocity than those produced by observers who experienced shorter delays. On average, observers whose lateral forces were measured 24 h after observation produced forces that were different from what they produced prior to force-field exposure; however, adapted force profiles for this group were variable, small in magnitude, and statistically unrelated to reach velocity. We obtained similar results from a control group that experienced the same 24 h delay period but did not observe force-field learning. This makes it difficult to attribute the effect we measured among observers who experienced a 24 h delay to force-field exposure. Thus, while our study found that the effects of observation on the human motor system can persist for at least 60 min after observing, we did not detect reliable evidence of observation-induced force-field adaptation 24 h after observation. In contrast, adaptation after physical force-field exposure was detectable at all time points probed, despite some decay after 24 h.

Previous studies have provided evidence that learning movement dynamics through observation involves some of the same cortical regions as learning through physical practice. Primary somatosensory (McGregor et al., 2016) and motor cortices (Brown et al., 2009) have been found to play necessary roles in acquiring new representations of reach dynamics, both for observing force-field learning and physical practice in a force-field. Regions of the cerebellum, intraparietal sulcus, and dorsal premotor cortex are also engaged both when physically experiencing reaching errors and when observing another individual undergoing force-field learning (Malfait et al., 2010). Observing force-field learning is also associated with changes in functional connectivity in a network involving the middle temporal visual area (V5/MT) and cerebellar, primary somatosensory, and primary motor cortices (McGregor and Gribble, 2015). This could provide a neuroanatomical basis for how visual information about observed movements reaches sensorimotor circuits to facilitate learning.

If observing and physically reaching in a force-field drive learning via common neural mechanisms, we might have expected to find that the passage of time equally affects the amount of adaptation. Instead, we found differences in how adaptation decayed after observation versus physical practice. Observation drove learning that was detectable for at least 1 h after the observation period, but which we did not detect after 24 h. In contrast, we could detect a signature of adaptation at all delay lengths after physical force-field exposure, with some decay after 24 h. Additionally, the temporal pattern of lateral forces was better preserved 1 h after physical force-field exposure compared to 1 h after observation. The adaptation induced by physically reaching in a force-field was therefore more temporally stable than that induced by observing force-field learning—a finding for which there may be at least two possible explanations.

First, it is possible that observation may drive adaptation via different neural mechanisms than learning through physical movement. This could lead to differences in the rates at which observation-and physical movement-based adaptation decay as time passes. A second option, which does not preclude the possibility that learning by observing and physical practice engage the same neural mechanisms, is that the differences we found in the temporal stability of adaptation could be attributable to differences in the amount of adaptation that occurred in the first place. The amount of adaptation induced by observation was about 30 % of that acquired through physical practice. It is possible that since observing drove less adaptation to begin with, it was also less stable over time. The idea that the initial amount of learning is related to its temporal stability is consistent with data previously reported by Joiner and Smith (2008), in which a longer training period and more complete adaptation resulted in increased retention of learning.

We also found differences in the rates at which observation-and physical practice-based adaptation washed out over repeated error clamp trials. Compared to movement training, observation-related learning washed out more slowly. While this difference could be interpreted as evidence of separate neural mechanisms, we cannot rule out the possibility that slower washout of observation-induced learning may instead reflect the fact that for observers, there was less adaptation available to be washed out in the first place. Only at the end of the washout period did levels of adaptation for participants who underwent physical training begin to approach the level of adaptation that was initially available to be washed out among observers. In studies of motor adaptation, washout following the removal of a perturbing stimulus characteristically follows a falling exponential curve (see Taylor and Ivry, 2014, for example). Washout therefore occurs at a faster rate in more highly adapted states, and at a slower rate in less adapted states. The washout rates reported for our observation and movement groups could therefore represent different phases of the same washout process, differentiated by the amount of adaptation left over at the time of the error-clamp probe.

### Potential Mechanisms Underlying Observational Learning

During motor learning through physical practice, changes in cortical representations of movement dynamics are thought to be driven by discrepancies between the sensed and predicted sensory consequences of motor commands, resulting in implicit adaptation (Wolpert and Miall, 1996; Shadmehr et al., 2010). During motor learning by observing, the brain learns about what forces are required for movement without the presence of efferent motor commands or afferent sensory feedback associated with one’s own movement. Instead, motor learning is driven by visual input.

It is possible that like physically reaching in a force-field, observing force-field learning can drive motor learning implicitly. In our study, observation-related adaptation of predictive limb control did not seem to depend upon observers’ conscious awareness of perturbing forces in the video they observed. We did not inform our participants about the presence of any external forces that may influence the reaching movements depicted in the video. Many participants in fact remarked when the video began to play that the tutor seemed to be “bad” at the task. This is consistent with participants attributing reaching errors to poor control on the part of the tutor rather than the presence of environmental disturbances. During informal discussions that occurred with a subset of participants after the experiment was completed (including more than half of all observers), observers were informed that the tutor was being perturbed by robot-induced forces. All said they had not previously been aware of any external forces acting on the arm they observed. Observers’ informal remarks during and after the experiment, taken together with the fact that they were instructed to follow the same rules as in the baseline block upon resuming reaching after observation, are consistent with the idea that observers likely did not adopt an explicit strategy to produce lateral forces in force channel probe trials following observation. This would also be consistent with the findings of Mattar and Gribble (2005) in which the effects of observation were not disrupted by a dual attention task engaging explicit cognitive systems.

The learning-related effects of observation on the motor system may be mediated by sensory prediction errors. Previous work has provided evidence that during action observation, observers use neural representations of movement to predict the actor’s movement kinematics. For example, Flanagan and Johansson (2003) showed that while observing a tutor perform a motor task, observers’ eye movements were predictively, rather than reactively, coordinated with movement of the tutor’s hand, suggesting that a feedforward process was used to predict the tutor’s hand kinematics and appropriately direct the observers’ gaze. In addition, a recent study by Kim et al. (2022) showed that observers learned to implicitly correct for errors they saw in an animation of a movement they had initially planned to execute but were cued to withhold, suggesting that prediction errors can drive adaptation even in the absence of movement. In the context of observation, it is possible that motor learning is driven in a similar way, using the discrepancy between visual information about the tutor’s reach kinematics and an observer’s implicit predictions of what reach kinematics ought to look like. Presumably in the present study such predictions would be informed by movement representations acquired during unperturbed reaching in the baseline block.

### Future Directions and Clinical Implications

Physical therapy to recover motor function after brain injuries and diseases affecting movement is commonplace. But for people affected by disorders that reduce their ability to undergo physical practice (for example, individuals affected by stroke with hemiparesis), an approach to neurorehabilitation that includes observation to aid in driving cortical reorganization may be a promising avenue for future research.

In the present study we have demonstrated that observing force-field learning induces motor adaptation and drives the formation of motor memories that can influence predictive limb control for at least an hour after the end of the observation period. The reorganization of motor circuits involved in force production and control has been recognized as a fundamental aspect of motor recovery after stroke (Kokotilo et al., 2009). Further, previous studies have provided evidence that motor cortical representations of dynamics for simpler tasks could contribute to representations of dynamics for more complex behavior (Gribble and Scott, 2002), providing a potential basis for how learning simpler motor skills could contribute to learning more complex ones. In our study, watching only a single, 12-minute video induced about 30 % of the adaptation gained through physically performing reaches in a novel dynamic environment. It could be that our video and observation protocol could be modified to produce stronger effects—for example, by increasing the size and number of reaching errors participants observe (Brown et al., 2010), or showing the observation video twice (Wanda et al., 2013). Carrying out multiple training sessions might also be helpful, since this has been shown to improve the learning and retention of skills learned through physical practice (Shea et al., 2000). If the differences we found in the temporal stability of observation-versus physical practice-induced adaptation were indeed attributable to differences in the amount of adaptation that occurred in the first place, then optimizing observational learning protocols to increase the amount of adaptation could also lead to the formation of more stable motor memories. It seems plausible that such interventions as those described above could improve the efficacy of observation-based interventions. However, whether such improvements would be enough for observation-based interventions to be clinically useful is a different question. Given that observation induced a rather small amount of adaptation in our study and in previous experiments (Mattar and Gribble, 2005; Wanda et al., 2013), we have doubts about the feasibility of using observation-based approaches alone to drive lasting cortical reorganization and clinically relevant behavioral change. A more fruitful alternative may be to leverage the effects of observation to enhance the effects of other interventions.

Observing force-field learning induces an adaptation of predictive limb control, which is thought to reflect reorganization in sensory and motor cortical regions. In our study, we found that this effect could be reliably detected for at least the first hour after observing. Future studies may explore whether observation-induced cortical reorganization could enhance the effects of physical interventions aimed at driving sensorimotor plasticity. We propose two broad mechanisms by which observing could augment the effects of physical practice.

First, if observation and physical practice were to contribute to the production and stabilization of the same representation of a skill, or even different representations that are both instantiated during skill execution, then perhaps it would be possible to use observation-based approaches to reduce the amount of physical therapy required for recovery of motor function after brain injury. Future studies may explore how much physical force-field reaching can be replaced by observation without reducing the total amount of learning that occurs. Further research may also explore the durability of the learning driven by combining physical practice and observation. For example, if participants were to learn 30 % of a skill by observing and then learn the remaining 70 % by physical practice, would the physical practice portion consolidate all of the learning that occurred?

Or would only the physical practice portion be retained, while the observation portion decays in a manner similar to what we report in our present study? Whether combining physical practice and observation produces effects that differ from the sum of their parts remains to be explored.

Second, it may be possible for observation to potentially augment the effects of subsequent physical practice by priming relevant neural circuitry. Previous studies provide evidence that regions that are critical for physical force-field learning (e.g., primary somatosensory cortex, primary motor cortex, and cerebellar regions) are also involved motor learning by observing (Brown et al., 2009; Malfait et al., 2010; McGregor et al., 2016; McGregor and Gribble, 2015). In inducing plasticity in those regions, one might expect that observation also induces the kinds of cellular changes (e.g. altered gene expression, changes in neuronal activity) that underlie reorganization (see Feldman, 2009, for review). Studies of visuomotor adaptation in which observation and physical practice are interleaved suggest that observation may replace some amount of physical practice Larssen et al. (2021), however the extent to which this may be true for motor learning tasks in which novel representations of movement dynamics are learned is unclear. Our findings suggest that observation-related changes in the neural circuitry underlying the acquisition and use of new representations of movement dynamics can persist for at least an hour after observing. This may provide a window of opportunity to enhance the potency of physical interventions that aim to drive plasticity in the same target regions.

